# ChromSCape : a Shiny/R application for interactive analysis of single-cell chromatin profiles

**DOI:** 10.1101/683037

**Authors:** Pacôme Prompsy, Pia Kirchmeier, Céline Vallot

**Affiliations:** CNRS UMR3244, Institut Curie, PSL Research University, 75005 Paris, France; Translational Research Department, Institut Curie, PSL Research University, 75005 Paris, France

## Abstract

Assessing chromatin profiles at single-cell resolution is now feasible thanks to recently published experimental methods such as single cell chromatin immunoprecipitation followed by sequencing (scChIP-seq) (Grosselin et al., 2019; Rotem et al., 2015) and single-cell assay for transposase-accessibility chromatin (scATAC-seq) (Buenrostro et al., 2015; Chen et al., 2018; Cusanovich et al., 2015; Lareau et al., 2019). With these methods, we can detect the heterogeneity of epigenomic profiles within complex biological samples. Yet, existing tools used to analyze bulk epigenomic experiments are not fit for the low coverage and sparsity of single-cell epigenomic datasets. Here, we present ChromSCape: a user-friendly Shiny/R application that processes single-cell epigenomic data to help the biological interpretation of epigenomic landscapes within cell populations. The user can identify different sub-populations within heterogeneous samples, find differentially enriched regions between subpopulations and identify associated genes and pathways. ChromSCape accepts multiple samples to allow comparisons of cell populations between and within samples. ChromSCape source code is written in Shiny/R, works as a stand-alone application and is freely downloadable at https://github.com/vallotlab/ChromSCape. Here, using ChromSCape on multiple H3K27me3 scChIP-seq datasets, we deconvolve chromatin landscapes within the tumor microenvironment, identifying distinct H3K27me3 landscapes associated to cell identity and tumor subtype.

## Introduction

Epigenetic landscape defined by histone modifications is driving chromatin folding and genes accessibility to transcription machineries. The recent development of single-cell methods to study the epigenome enables the appreciation of the heterogeneity of chromatin features within a population, which cannot be assessed using bulk ATAC-seq or ChIP-seq method. Such methods include scChIP-seq (Grosselin et al., 2019; Rotem et al., 2015), scChIL-seq (Harada et al., 2019), scChIC-seq (Ku et al., 2019) and scCUT&Tag (Kaya-Okur et al., 2019) which identify genomic regions enriched for repressive or active histone marks (H3K27me3, H3K4me3, …) and scATAC-seq (Chen et al., 2018; Cusanovich et al., 2015; Lareau et al., 2019) which assesses regions of open chromatin. There are existing tools publicly available to analyze scATAC-seq such as chromVar (Schep et al., 2017), Cicero (Pliner et al., 2018) or Scasat (Baker et al., 2019). However these tools are dedicated to scATAC-seq and are not stand-alone applications, thus requiring some scripting skills. To the best of our knowledge, there has not been any stand-alone analytic tool destined to comprehensively analyze region-based scChIP-seq count data. With ChromSCape, we propose a user-friendly, step-by-step and customizable Shiny application to analyze sparse single-cell chromatin profiling datasets. The pipeline is designed for high-throughput single-cell datasets with samples containing as low as 100 cells and with a minimum of 1000 reads per cell. The interactive process includes filtering out cells with low coverage and regions with low cell count, dimensionality reduction by PCA, classifying cells in an unsupervised manner to identify sub-populations and find biologically relevant loci differentially enriched in each sub-populations. To overcome the sparsity of matrices due to the current technical limitations of these technologies, reads mapped to the genome must be binned prior to the use of ChromSCape into successive bins of the genome. These bins range from 5,000bp (recommendation for H3K4me3 scChIP-seq datasets) up to 50,000bp (recommendation for H3K27me3 scChIP-seq datasets), depending on the modification studied, to increase signal with a resolution trade-off.

## Implementation

ChromSCape is developed in Shiny/R employing various Shiny related packages (shinyjs, shinydashboard, shinyDirectoryInput) for the user interface. The application takes advantage of public R libraries for data vizualisation (RcolorBrewer, colorRamps, Rtsne, colourpicker, kableExtra, knitr, viridis, ggplot2, gplots, png, grid, gridExtra, DT) as well as for data manipulation (tibble, dplyr, tidyr, stringr, irlba, reshape2, splitstackshape, rlist). Some available bioinformatics packages are used for manipulation of single-cell data (scater (McCarthy et al., 2017), scran (Lun ATL and JC, 2016)), manipulation of genomic regions (IRanges and GRanges (Lawrence et al., 2013)) and for clustering of cells (ConsensusClusterPlus (Wilkerson et al., 2010)). Some custom functions are embedded in the application under the ‘Modules’ directory and serve for both manipulation and vizualisation of data sets. Brief command lines provided in https://github.com/vallotlab/ChromSCape enable users without any informatic background to install all R dependencies and run the application in a web browser. A run command-line program achieving similar results is also available at https://github.com/vallotlab/scChIPseq.

## Methods

ChromSCape consists of multiple filtering and processing steps, saving files on a user specified directory at each step. When the user relaunches ChromSCape and selects a saved analysis, the application automatically reloads the data and recomputes all the plots. This enables users to try various filtering and clustering parameters, visualize the data in reduced dimensional space and choose the most appropriate set of parameters for their analysis.

### Quality Control and Normalization

ChromSCape takes as input one or multiple count matrices with genomic regions in rows and cells in columns. Each input matrix should have the same regions as rows to allow merging of the different matrices. An example matrix is available on the GitHub repository; guidelines given within the application allow users to understand the requirements for input formating. In order to efficiently remove outlier cells from the analysis, e.g. cells with excessively high or low coverage, the user sets a threshold on a minimum read count per cell and the upper percentile of cells to remove. The latter could correspond to doublets, e.g. two cells in one droplet, while lowly covered cells are not informative enough or may correspond to barcodes ligated to contaminant DNA or library artifacts. Regions not supported by a minimum user-defined percentage of cells that have a coverage greater than 1,000 reads are filtered. Defaults parameters were chosen based on the analysis of multiple scChIP-seq datasets from our previous study (Grosselin et al., 2019): a minimum coverage of 1,600 unique reads per cell, filtering out the cells with the top 5% coverage and keeping regions detected in at least 1 % of cells. Post quality control filtering, the matrices are normalized by dividing each cell by its total read count, and multiplying by the average total read count across all cells. At this step, the user can provide a list of genomic regions, in BED format, to exclude from the subsequent analysis, in cases of known copy number variation regions between cells for example.

### Dimensionality Reduction

In order to reduce the dimensions of the normalized matrix for further analysis, principle component analysis (PCA) is applied to the matrix, with centering, and the 50 first PCs are kept for further analysis. The user can visualize scChIP-seq data after quality control in the PCs dimensional space. The t-distributed stochastic neighbor embedding (t-SNE) algorithm (Maaten and Hinton, 2008) is applied on the PCA to visualize the data in two dimensions. The PCA and t-SNE plots are a convenient way to check if cells form clusters in a way that was expected before any clustering method is applied. For instance, the user should verify whether the QC filtering steps and normalization procedure were efficient by checking the distribution of cells in PC1 and PC2 space. Cells should group independently of normalized coverage. In our hands, for our scChIP-seq H3K27me3 datasets, minimum coverage of 1,600 unique reads per cell was required to separate cells independently of coverage post normalization (Grosselin et al., 2019). We haven’t observed any batch effect between our experiments in our first study most probably because we were working with a single batch of hydrogel beads. Therefore no correction is implemented in this version of the application, but we are planning to implement solutions to correct batch effect in a future version.

### Correlation and Filtering

Using the normalized dataset as input, hierarchical clustering is performed on the pairwise Pearson’s correlation matrix. To improve the stability of our clustering approaches and to remove from the analysis isolated cells that do not belong to any subgroup, cells displaying a Pearson’s pairwise correlation score below a threshold *t* with at least *p* % of cells are filtered (*p* is recommanded to be set at 1 % or 2 % depending on the dataset). The correlation threshold *t* is calculated as a user-defined percentile of Pearson’s pairwise correlation scores for a randomized dataset (percentile is recomanded to be set as the 99th percentile). The correlation heatmaps before and after correlation filtering and the number of remaining cells are displayed to inform users on the filtering process.

### Consensus Correlation Clustering

ChromSCape uses Bioconductor ConsensusClusterPlus package (Wilkerson et al., 2010) to examine the stability of the clusters and compute item consensus score for each cell. Consensus partitions of the dataset into a number of cluster ranging from 2 to 10 is done on the basis of 1,000 resampling iterations (80% of cells sampled at each iteration) of hierarchical clustering, with Pearson’s dissimilarity as the distance metric and Ward’s method for linkage analysis. The optimal number of clusters can be chosen in order to maximize intra-cluster correlation scores based on the graphics displayed on the ‘Consensus Clustering’ tab after processing. Clustering results can also be visualized in two dimensions with the t-SNE plot. This unsupervised clustering allows to discover clusters of cells based on their respective chromatin profile within a population without any prior knowledge.

### Peak Calling for genomic region annotation

This step of the analysis is optional, but recommended in order to obtain meaningful results with the enrichment analysis. To be able to run this module, some additional command line tools are required such as Samtools (Li et al., 2009), Bedtools (Quinlan and Hall, 2010) and MACS2 (Liu, 2016). The user needs to input BAM files for the samples (one separate BAM file per sample), with each read being labeled with the barcode ID. ChromSCape merges all files according and split them again according to the previously determined clusters of cells (one separate BAM file per cluster). Customizable significance threshold for peak detection and merging distance for peaks (defaults to p-value=0.05 and peak merge distance to 5,000) allows to identify peaks in close proximity (<1000bp) to a gene transcription start site (TSS); these genes will be later used as input for the enrichment analysis. For the annotation, ChromSCape uses the reference human transcriptome Gencode hg38 v26 (Frankish et al., 2018), limited to protein coding, antisense and lncRNA genes.

### Differential Analysis and Gene Set Enrichment Analysis

To identify differentially enriched regions across single-cells for a given cluster, ChromSCape performs a non-parametric two-sided Wilcoxon rank sum test comparing normalized counts from individual cells from one cluster versus all other cells. We test for the null hypothesis that the distribution of normalized counts from the two compared groups have the same median, with a confidence interval 0.95. The calculated p-values are then corrected by the Benjamini-Hocheberg procedure (Benjamini and Hochberg, 1995). The user can set a log2 fold-change threshold and corrected p-value threshold for regions to be considered as significantly differentially enriched (we recommend to set corrected p-value and log2 fold-change thresholds respectively to 0.01 and 1). Using the refined annotation of peaks done in previous step, the final step is to look for enriched gene sets of the MSigDB v5 database (Subramanian et al., 2005) in differentially enriched regions. We apply hypergeometric tests to identify gene sets from the MSigDB v5 database overrepresented within differentially enriched regions, correcting for multiple testing with the Benjamini-Hochberg procedure. Users can then visualize most significantly enriched or depleted gene sets corresponding to the epigenetic signatures of each cluster and download gene sets enrichment tables.

## Application : ChromSCape deconvolves chromatin landscapes of the tumor micro-environment

To showcase the use of ChromSCape to interrogate heterogeneity of chromatin landscapes within several samples, we analyzed together four H3K27me3 mouse scChIP-seq datasets (GSE117309), two of which had not been analyzed in our previous study (Grosselin et al., 2019). The raw FASTQ reads were processed using the latest version of our scChIP-seq data engineering pipeline that allowed a more precise removal of PCR and RT duplicates (code available at <u>Github</u>). Count matrices (available at <u>Figshare</u>) were formatted into a ChromSCape compliant format. The samples correspond to the mouse cells from patient-derived xenograft (PDX) originating from two different human donors. The HBCx-22 and HBCx-22-TamR datasets correspond to mouse cells from a pair of luminal ER^+^ breast PDXs: HBCx-22, responsive to Tamoxifen and HBCx-22-TamR, resistant to Tamoxifen. The HBCx-95 and HBCx-95-CapaR correspond to a triple-negative breast cancer (TNBC) tumor model of acquired resistance to chemotherapy. The four 50,000-bp binned matrices of raw counts, in mm10 reference genome, were given as input to ChromSCape to interrogate the heterogeneity of chromatin states within the tumor micro-environment of both luminal and TNBC tumors. Tumor micro-environment is a key player in tumor evolution processes, and can vary between tumor types and with response or resistance to cancer therapy. With ChromSCape, we propose a comprehensive view of cell populations based on their chromatin profiles, and the identification of tumor-type and treatment-specific cell populations and respective chromatin features. All figures excepted Fig. 2d were automatically produced by the application and are easily downloadable.

**Fig. 1.**
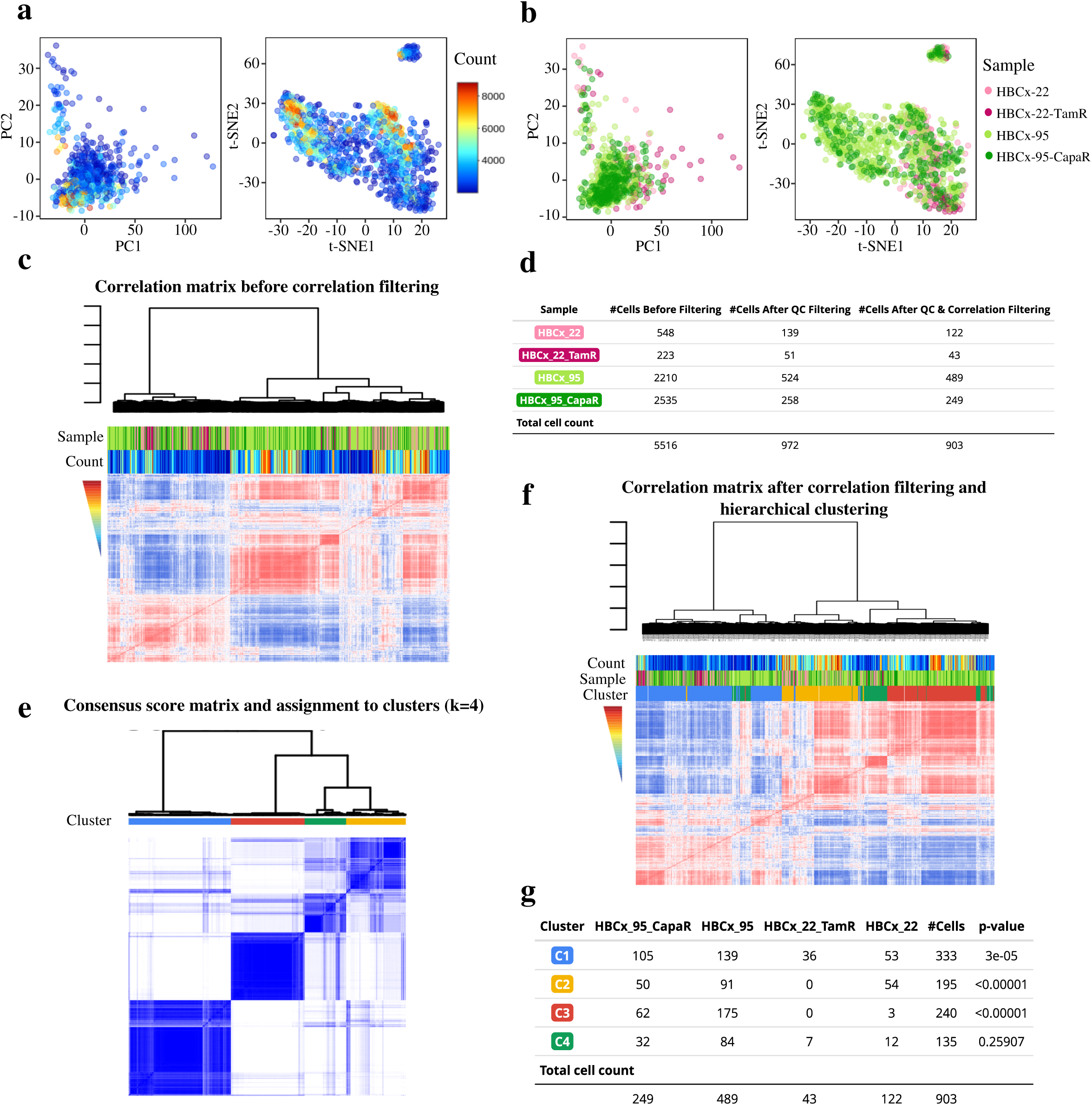
Filtering and Unsupervised Clustering of single-cell ChIP-seq H3K27me3 histone mark profiles of mouse stromal cells from patient derived xenografts models. (HBCx-22, HBCx-22-TamR, HBCx-95, HBCx-95-CapaR PDXs, see Grosselin et al., 2019). Cells with lower than 2000 counts or with higher counts than the 95th percentile were filtered. Regions not represented in at least 1% (*n*=23) of remaining cells were excluded. (a) PCA and t-SNE plots colored by unique mapped reads. (b) PCA and t-SNE plots colored by sample of origin. (c) Hierarchical clustering of cell-to-cell Pearson’s correlation scores before filtering step. (d) Table of each sample’s cell number before and after correlation filtering. (e) Hierarchical clustering and corresponding heatmap of cell-to-cell consensus clustering of cells using *k* = 4 clusters. Consensus score ranges from 0 (white: never clustered together) to 1 (dark blue: always clustered together). Cluster membership is color coded above the heatmap. (f) Hierarchical clustering and corresponding heatmap of cell-to-cell Pearson’s correlation scores for ‘correlated’ cells only. (g) Table of samples memberships to the clusters. P-value column results from Pearson’s Chi-squared goodness of fit test without correction, checking if the observed distribution of samples in each cluster differs from theoretical distribution.

**Fig. 2.**
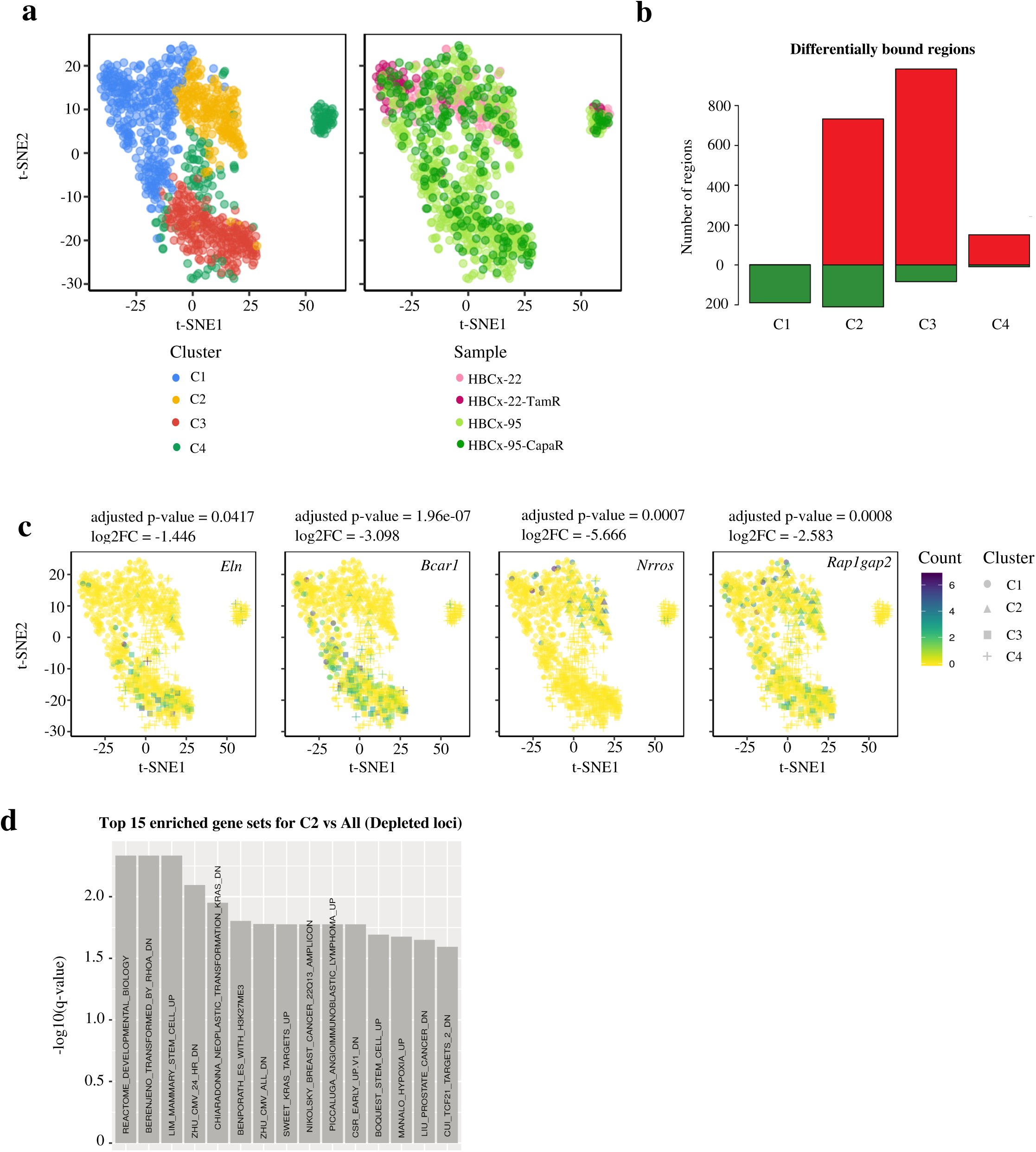
Differential Analysis and Gene Set Enrichment Analysis of single-cell ChIP-seq H3K27me3 histone mark profiles of mouse stromal cells. (samples HBCx-22, HBCx-22-TamR, HBCx-95, HBCx-95-CapaR PDXs). (a) t-SNE representations after correlation filtering (n=903 cells) and colored by cluster or sample of origin. (b) Differentially bound regions identified by Wilcoxon signed-rank test. Genomic regions were considered enriched (red) or depleted (green) in H3K27me3 if the adjusted p-values were lower than 0.01 and the fold change greater than 2. (c) t-SNE representation of scChIP-seq dataset, points are colored according to H3K27me3 enrichment signals in each cell for genes located in depleted regions in C1 to C4, respectively *Eln, Bcar1, Nrros* and *Rap1gap2*. The adjusted p-values and log2FC of the associated regions are indicated above each plot. (d) Barplot displaying the −log10 of adjusted p-values from pathway analysis for cells of cluster C2 compared to all other cells in depleted loci. Only the top 15 significant gene sets are indicated, filtering out depleted gene sets from MSIGdb classes “c2_computational” and “c3_motifs”.

In the quality filtering step, a threshold of 2,000 minimum reads per cell was set due to a relatively high initial number of cells (5,516 cells in total, see Fig. 1d). Samples were not affected to the same extent by the filtering step (Fig. 1d, p-value = 5e-04, Fisher’s exact test): sample HBCx-95-CapaR was affected twice more than all other samples by the filtering probably due to a lower initial cell coverage. After normalization and dimensionality reduction, the data is visualized in reduced dimensional spaces (after PCA and t-SNE reduction steps) according to either total count or sample of origin (Fig. 1a-b). In both representations, we can observe multiple clouds of high count cells, showing that total count is not the only source of variation between cells post normalization. Interestingly, cells from the two different model mix generally well together showing that no major batch effect was present between samples. Correlation filtering (Fig. 1c-d) was done setting the quantile threshold at 99%, *i.e* cells need to have a correlation score greater than 0.44 with at least 1% of other cells to be kept for downstream analysis (Sup. Fig. 1). We note that this filtering step may be biased towards removing meaningful cells from samples with low cell number, as they might not have enough representative cells with each chromatin profile to pass treshold.

After performing consensus clustering approach on the filtered dataset for *k*=2 to *k*=10 clusters, we chose to partition the data into *k*=4 clusters based on the knee method as a major leap in area under the CDF curve was observed between *k*=3 and *k*=4 clusters, and not between *k*=4 and *k*=5 (see Sup. Fig. 2 a-b). Consensus score matrix in Fig. 1e shows that most of the cells were stably assigned to four chromatin-based populations. Cluster C2 and C4 cells assignment is less stable than C1 and C3 (mean consensus scores are respectively 0.84 and 0.90 for C1-C3 and 0.70 and 0.71 for C2-C4, see Sup. Fig. 2.a), suggesting that cells from C2 and C4 might share H3K27me3 features, whereas cells from C1 and C3 have distinct H3K27me3 landscapes. Clusters C1, C2 and C4 contain cells from all four samples, with a significantly higher proportion of HBCx-22-TamR for C1 (p-value = 3e-05, Pearson’s Chi-squared test) (Fig. 1f, 1g, 2a). On the other hand, cluster C3 is almost exclusively (p-value < 1e-05) composed of cells from model HBCx-95 (Fig. 1b, 2a), revealing a stromal cell population specific to the triple negative breast cancer model (HBCx-95).

To further identify the specific features of each chromatin-based population, we proceeded to peak calling, differential analysis and gene set enrichment analysis using default parameters (see Methods). As H3K27me3 is a repressive histone mark, we focused our analysis on loci depleted in H3K27me3, where transcription of genes can occur. The differential analysis identified respectively 189, 210, 83 and 9 depleted regions for clusters C1 to C4 (Fig. 2b). We found loci devoid of H3K27me3 specific to cluster C2, enriched for genes involved in apical junction such as *Bcar1* (Fig. 2c) and *Ptk2*, which are characteristic of genes expressed in fibroblasts. We found a depletion of H3K27me3 specific to cluster C3 over the genes *Nrros* (Fig. 2c) and *Il10ra*, two genes characteristic of immune expression programs. Depletion of H3K27me3 over the transcription start site of *Rap1gap2*, a gene expressed in endothelial cells, was a key feature of cluster C4 (Fig. 2c). For cluster C1 and C2, we found a depletion of H3K27me3 over *Eln*, a gene expressed in fibroblasts.

Gene set enrichment analysis (q-value < 0.1) for genes located in regions depleted of H3K27me3 enrichment only revealed very few enriched gene lists, mostly for cluster C2 (Fig. 2d, multiple gene sets related to stem and cancer cells) and one list for C1 (“LPS_VS_CONTROL_MONOCYTE_UP”) and (Fig. 2b). Linking H3K27me3 enrichment to transcription is indeed indirect, we see our gene list tool more appropriate for H3K4me3 scChIP-seq or scATAC-seq datasets, where enriched regions are directly associated to gene transcription. In addition, we are working on improving this step in future versions of the application, including enhancer annotation for example.

Overall, these results are consistent with our previous analysis of HBCx-95 scRNA-seq datasets where subpopulations were differentially expressing markers of fibroblasts, endothelial and macrophage cells (Grosselin et al., 2019). This new analysis comprising the HBCx-22 dataset allowed us to identify the H3K27me3 signature of potential endothelial cells (cluster C4). These cells are present in each model, but might not have been previously detected in the previous scChIP-seq analysis due to low cell representation. In addition, the H3K27me3 signature of potential immune cells is restricted to cells from the TNBC model (cluster C3), suggesting that these immune cells are absent from the luminal tumor. Altogether, ChromSCape can effectively deconvolve chromatin landscapes in complex samples, such as tumors.

## Conclusion

ChromSCape is a standalone Shiny/R application designed for both biologists and bioinformaticians to analyze complex chromatin profiling datasets such as scChIP-seq datasets. The comprehensive application is quick to take over plus the direct visualization of cells clusters combined to configurable parameters and incremental saving of intermediary R objects eases bench-marking of parameters. With an analysis of several scChIP-seq datasets of mouse stromal cells, we show that heterogeneity of chromatin landscapes between and within cell populations can robustly be detected by implementing multiple filtering steps as well as consensus clustering approach. Overall, we predict that ChromSCape will be a useful tool to probe heterogeneity and dynamics of chromatin profiles in various biological settings, not only in cancer development but also in cell development and cellular differentiation.

---

**Supplementary Fig. 1.**
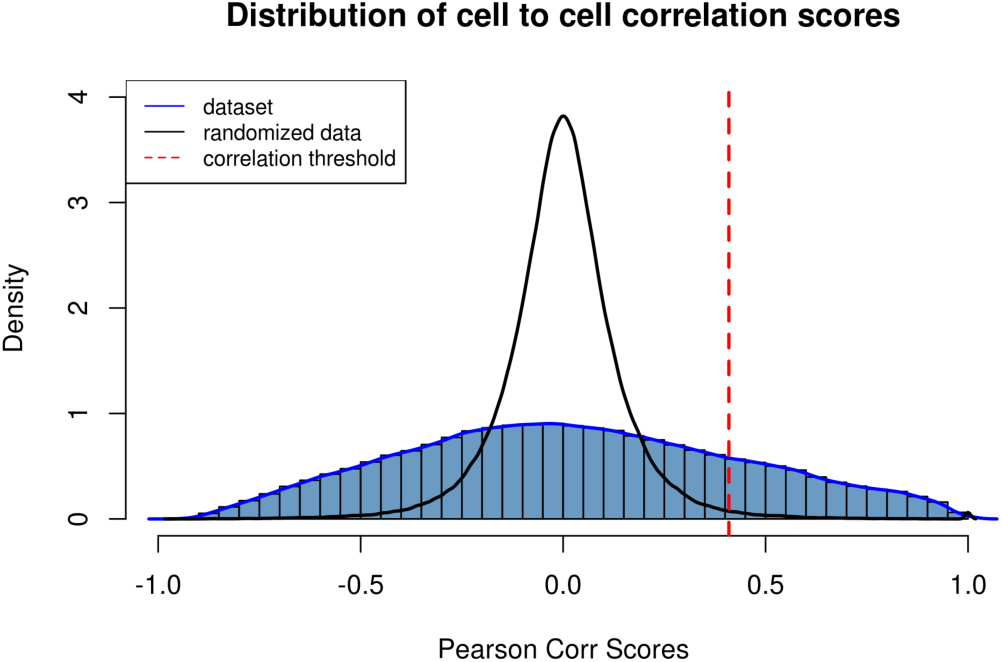
Distribution of cell to cell correlation scores. Correlation threshold is calculated as the 99th quantile of the randomized dataset (*t=*0.44).

**Supplementary Fig. 2.**
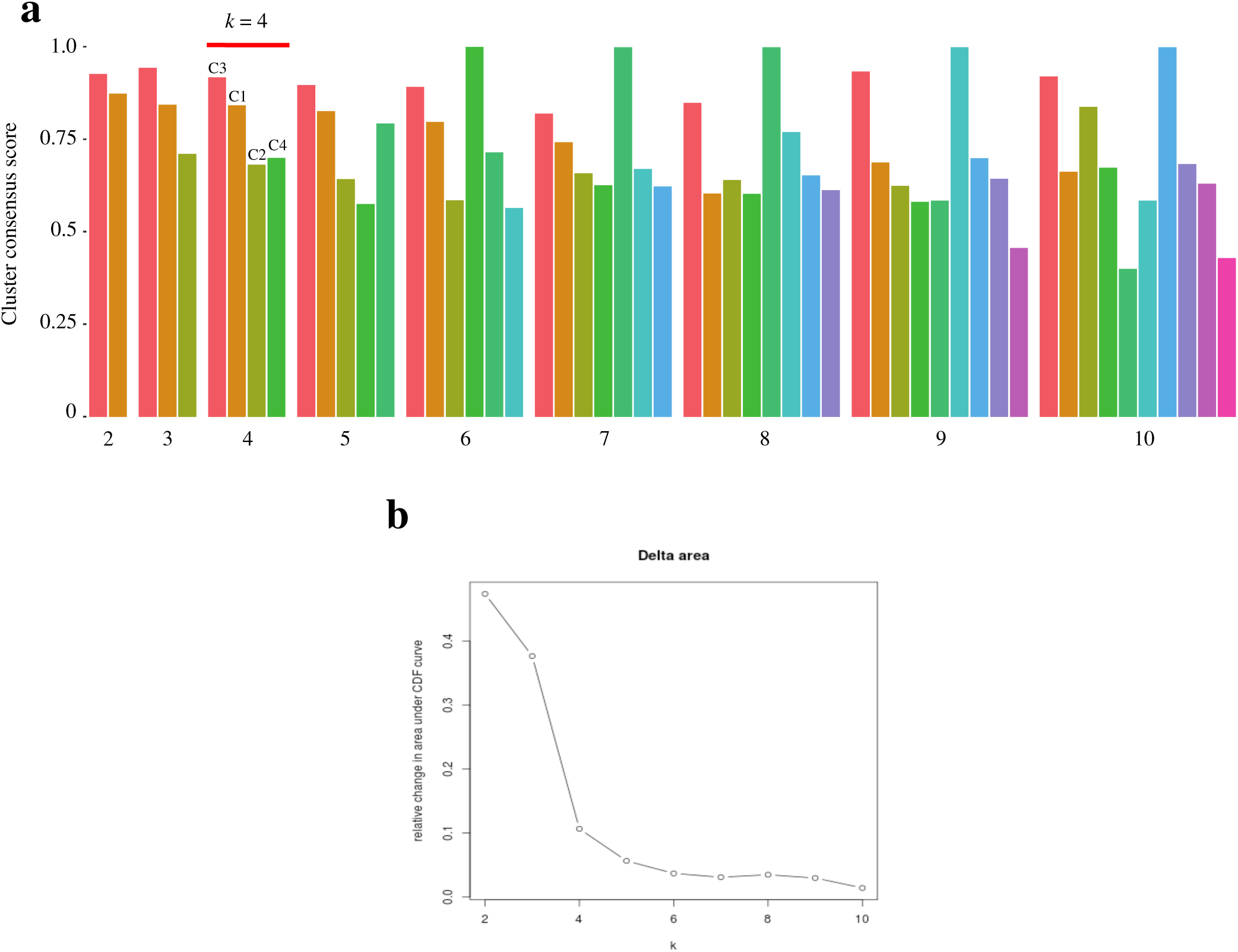
Consensus clustering of n=903 mouse stromal cells using 80% of cells at each iteration and 1,000 iterations. (samples HBCx-22, HBCx-22-TamR, HBCx-95, HBCx-95-CapaR PDXs). (a) Barplot of consensus scores for each segmentation, from k=2 to 10. *k=*4 clusters was chosen. The names of the clusters corresponding to the analysis is showed above histogram for *k*=4. (b) Relative change in area under the Cumulative Distribution Fraction for k=2 to 10 clusters.

